# Surface Functionalized RBC Membrane-Derived Nanoparticles for Targeted Drug Delivery to Attenuate Fatty Liver Disease

**DOI:** 10.64898/2026.02.23.707593

**Authors:** Alap Ali Zahid, Jiaqi Huang, Nica Borradaile, Arghya Paul

**Affiliations:** Department of Chemical and Biochemical Engineering, The University of Western Ontario, London, ON N6A 5B9, Canada; Department of Physiology and Pharmacology, Schulich School of Medicine and Dentistry, Western University, London, Ontario N6A 3K7, Canada; Department of Chemistry, The University of Western Ontario, London, ON N6A 5B9, Canada; Collaborative Specialization in Musculoskeletal Health Research and Bone and Joint Institute, The University of Western Ontario, London, ON N6A 5B9, Canada; School of Biomedical Engineering, The University of Western Ontario, London, ON N6A 5B9, Canada

**Keywords:** Cell membrane-derived nanoparticles, RBCs, drug delivery, self-assembled nanoparticles, MASLD, hepatic steatosis

## Abstract

Metabolic dysfunction-associated steatotic liver disease (MASLD) is marked by excessive hepatic lipid accumulation and is closely associated with hyperlipidemia. It poses significant health challenges and can progress to severe chronic liver disease if untreated. Several small-molecule pharmacological agents are either in clinical use (resmetirom) or advancing through preclinical development for the treatment of hepatic steatosis. However, some promising lead drug candidates have limited therapeutic potential due to poor solubility, low permeability, limited biocompatibility, and off-target effects. Cell membrane-derived nanoparticles (CMN), prepared from red blood cells, naturally exhibit immune-evasion properties and can overcome these limitations by encapsulating small molecules within their self-assembled structures. Further, CMN can be surface functionalized to enable precise targeting of liver hepatocytes. Here, we developed a hepatocyte-targeting CMN loaded with a model drug (resmetirom) for MASLD therapy. Using covalent bonds, we conjugated three different hepatocyte-targeting ligands to CMN and identified lactoferrin as the most effective ligand through comparative screening. We then confirmed the cellular internalization pathways of the selected ligand in both targeted CMN and non-functionalized CMN. Finally, in an *in vitro* hepatic steatosis model, the optimized targeted CMN demonstrated improved bioactivity, including significant reductions in lipid droplets, triglycerides, and liver enzyme levels. Altogether, this targeted CMN platform shows promising potential to enhance the therapeutic efficacy of small-molecule drugs for MASLD and may, overall, improve therapeutic outcomes in preclinical and clinical trials.

## 1. Introduction

Approximately one-third of the world’s population suffers from metabolic dysfunction-associated steatotic liver disease (MASLD). MASLD is a complex, progressive metabolic disorder that begins with excess fat accumulation in the liver, progresses to metabolic dysfunction-associated steatohepatitis (MASH), and in later stages can lead to liver failure or hepatocellular carcinoma.^1,2^ Fat accumulation exceeding 5% in hepatocytes is considered a hallmark of MASLD.^3,4^ In addition, MASLD commonly coexists with hypertriglyceridemia, hypercholesterolemia, obesity, diabetes, and hypertension. Projections indicate that by 2030, cases of advanced liver disease and liver-related deaths due to MASLD will more than double, presenting an urgent and significant global healthcare challenge.^3,5,6^ Most drugs indirectly treat the disease by alleviating its progression, such as treating type 2 diabetes mellitus, reducing excess cholesterol in the bloodstream, and improving oxidative stress.^7–9^ However, the overall efficacy of such pharmacotherapy for MASLD remains poor. Recently, the FDA approved GLP-1 receptor agonists such as Semaglutide for MASH therapy.^10,11^ Moreover, Rezdiffra (resmetirom) was FDA-approved for the treatment of MASLD-associated fibrosis and is currently available in the US as a first-line therapeutic option.^12,13^ During the phase III clinical trial of resmetirom (Res), 25.9% and 29.9% of patients treated with 80 mg and 100 mg resmetirom, respectively, achieved MASLD/MASH resolution. ^13^ Ideally, compared with oral administration of Res, intravenous administration could enhance hepatic targeting and bioavailability by bypassing gastrointestinal degradation, but its poor water solubility limits its clinical use.^11,14^Collectively, these agents demonstrate meaningful progress in MASLD/MASH therapy; however, adverse events, including severe diarrhea and nausea, were also observed in patients during the trial. However, there is potential to improve the therapeutic efficacy of Res, a BCS (Biopharmaceutics Classification System) Class IV drug, through advanced formulations. ^15,16^

New techniques using synthetic nanoparticles (e.g., liposomes), exosomes, and extracellular vesicle-based drug-delivery systems have been explored to enhance the delivery of therapeutics and thereby improve their efficacy. However, synthetic nanoparticles face several limitations, including poor biocompatibility, reduced circulation time, limited cellular uptake, and a lack of targeting.^17–19^ Similarly, exosome- and extracellular vesicle-based systems are limited by low production yields, time-consuming preparation, and the requirement for large numbers of source cells.^20^ Hence, there is an urgent need to develop an efficient drug-delivery platform to enhance the efficacy of drugs for the treatment of MASLD. Among these nanoparticles, we propose red blood cell (RBC) membrane-derived nanoparticles (CMN), prepared directly from natural RBC membranes, as an effective drug-delivery platform that improves biocompatibility, minimizes systemic toxicity, and enhances delivery efficiency to liver cells. ^21–23^ These RBC-sourced CMN can be generated from autologous cells or from patient-matched blood obtained from healthy donors, thereby broadening the scope for personalized, biocompatible, and clinically translatable nanotherapeutic strategies.^17,24^ Furthermore, the RBC membrane expresses immune-regulatory proteins, such as CD47, which sends a “don’t eat me” signal that inhibits macrophage phagocytosis, along with the presence of abundant sialylated glycans that prolong circulation and reduce immune clearance.^25,26^

Separately, the liver is a metabolic center that collects, metabolizes, and transports substances throughout the body, and is mainly composed of hepatocytes (parenchymal cells). Delivering and retaining drugs in hepatocytes is crucial because MASLD and other chronic liver diseases originate in these cells.^1,27^ However, when nanoparticle-based systems are used, they are often taken up by phagocytes of the reticuloendothelial system (RES), particularly Kupffer cells, which are non-parenchymal liver cells. Therefore, to improve the effectiveness of CMN-based nanocarriers, it is essential to functionalize their surfaces with specific ligands that enable active targeting and binding to hepatocyte receptors.^28,29^ This receptor-mediated endocytosis provides a promising strategy for delivering drugs to hepatocytes, resulting in high intracellular uptake while minimizing off-target exposure. The asialoglycoprotein receptor (ASGPR) is abundantly expressed on hepatocytes and minimally expressed in extrahepatic tissues, offering a significant advantage for targeted delivery to hepatocytes.^30,31^ ASGPR is well known for its cell internalization by clathrin-mediated endocytosis and shows high affinity for galactose, carbohydrates, and glucose.^30^ Furthermore, sugar isomers, galactose density and branching, spatial configuration, and galactose linkages all play critical roles in ligand-receptor binding. Multiple distinct ligands have been reported for receptor-mediated targeted delivery of a drug to the liver, identifying an optimal ligand with high affinity for hepatocytes is much needed.

In this study, we selected three targeting ligands and screened for the one with the highest hepatocyte binding, as illustrated in **Figure 1**. CMN can be readily surface functionalized with targeting molecules via amine groups, facilitating efficient surface modification. We selected glycyrrhetinic acid (Ga), lactobionic acid (La), and lactoferrin (Lf) as targeting ligands for this study. Ga is one of the main bioactive compounds in licorice and is a potential ligand for glycyrrhetinic acid receptor and ASGPR on hepatocytes.^32,33^ Similarly, La is known for its effective binding to the ASGPR, and researchers have been using it for active targeting in drug delivery to alleviate fatty liver disease.^29^ Lastly, Lf, a cationic glycoprotein, has been shown to bind multiple receptors on hepatocytes and has become increasingly attractive for its high-affinity binding to ASGPR.^34^ We screened CMN-Ga, CMN-La, and CMN-Lf for enhanced cell internalization into hepatocytes using flow cytometry. Once the optimal CMN-based targeting ligand was identified, we assessed bioactivity by loading the model drug (Res) into the functionalized CMN and comparing Res-loaded functionalized CMN efficacy with that of the free drug in an *in vitro* model of steatotic hepatocytes. This work supports the concept that CMN-based nanocarriers have significant potential as targeted drug-delivery platforms, enabling the delivery of pharmacological agents or other small molecules directly to hepatocytes, and may represent an effective strategy for alleviating MASLD.

**Figure 1.**
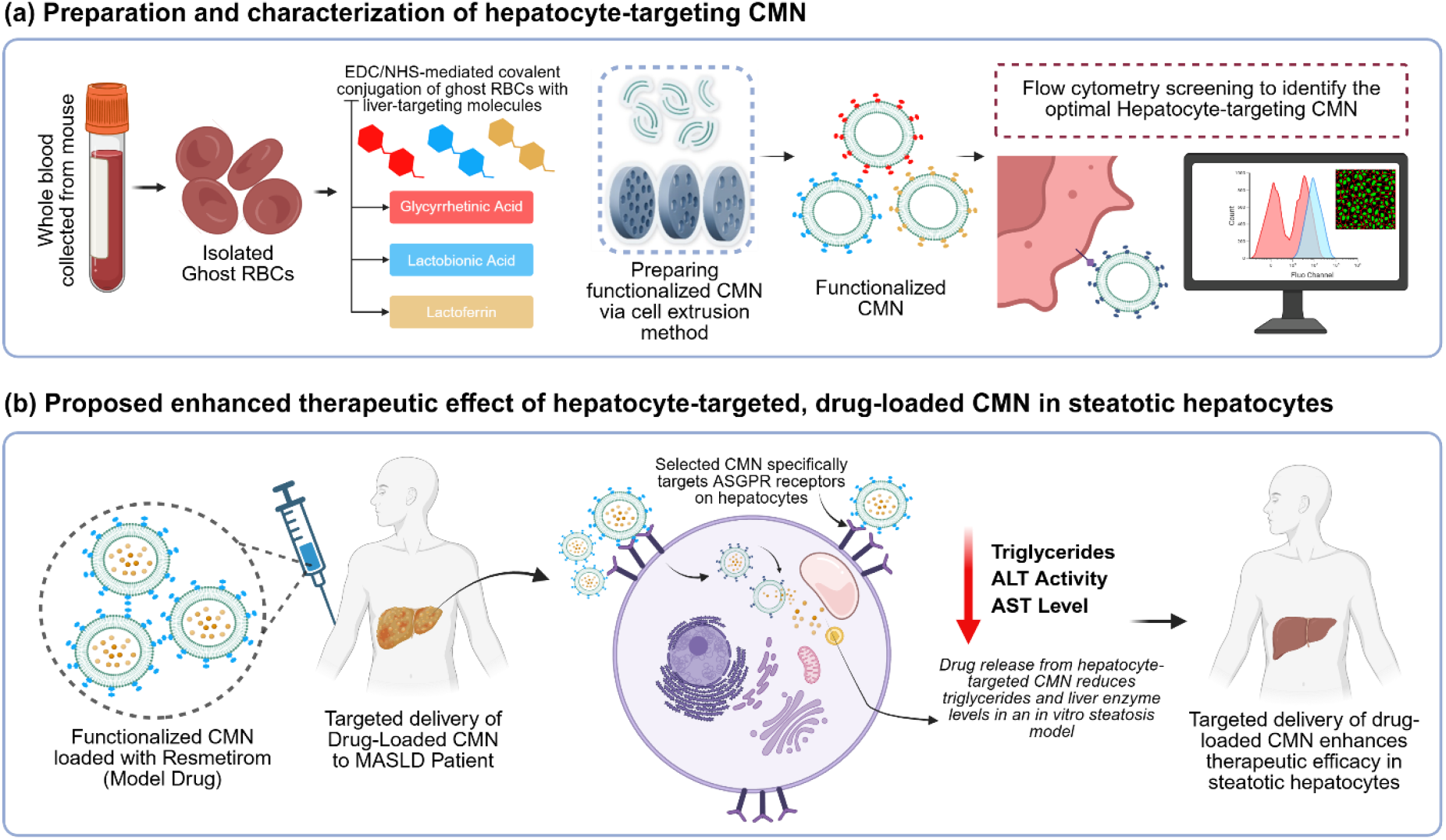
Schematic illustration of proposed hepatocyte-targeting CMN derived from RBCs to improve the therapeutic efficacy of the model drug (resmetirom) in steatotic hepatocytes. a) Preparation and characterization of functionalized RBC-sourced CMN surface-modified with glycyrrhetinic acid, lactobionic acid, and lactoferrin, followed by flow cytometry screening to identify the optimal hepatocyte-targeting formulation using a cellular uptake assay. b) The selected hepatocyte-specific CMN loaded with resmetirom demonstrates improved efficacy by reducing intracellular triglycerides and lowering elevated liver enzyme levels in steatotic hepatocytes.

## 2. Experimental section

### 2.1. Materials

Human HepG2 cells (Hep G2-HB-8065, ATCC) at passage 20 were cultured in Eagle Minimum Essential Medium (EMEM; M5650, Millipore-Sigma). Complete media for culturing HepG2 cells was prepared by adding 10% (v/v) fetal bovine serum (FBS) (Gibco, TMS-016-B), 1% (v/v) penicillin-streptomycin (Gibco, 15140122), and 2 mM L-Glutamine (Millipore-Sigma, 59202C). THP-1, a human monocytic leukemia cell line, was cultured in RPMI 1640 medium containing 10% (v/v) FBS, 2mM L-glutamine at 37 °C under 5% CO2. A live cell tracker or DiI stain for tagging the CMN was purchased from Millipore Sigma (42364). The cell apoptosis kit (Annexin V and propidium iodide, PI) was purchased from Biotium (30061). Cell mitochondrial changes and cell viability were assessed using the MTS assay (Promega, G3580) and live/dead cell imaging kit (LIVE/DEAD™ Cell Imaging Kit, R37601), respectively. All other chemicals, including 18β-Glycyrrhetinic acid (G10105), lactobionic acid (L9507), lactoferrin from bovine milk (L9507), N-Hydroxysuccinimide/NHS (130672), and N-(3-Dimethylaminopropyl)-N′-ethylcarbodiimide hydrochloride/EDC (25952-53-8), and Oil Red O (1320-06-5) were purchased from Millipore-Sigma, Canada. Triglycerides, alanine aminotransferase (ALT), and aspartate aminotransferase (AST) kits were purchased from Cayman, with catalogue numbers 10010303, 701640, and 700260, respectively. Whole mouse blood (SKU: IGMSCD1WBNAC10ML) was purchased from Innovative Research, USA.

### 2.2. Preparation of hepatocyte-targeted CMN using erythrocyte (RBC) ghost membranes

Ghost RBC membrane-derived CMN were prepared according to the previously published protocol.^21,26^ In brief, whole mouse blood was used to isolate ghost RBCs after hypotonic treatment, and isolated ghost RBCs were quantified using a hemocytometer. Next, the quantified RBCs were used to prepare CMN and liver targeting CMN functionalized with glycyrrhetinic acid (Ga), lactobionic acid (La), and lactoferrin (Lf), as further explained in the following subsections.

#### Ga-modified RBCs

Ga was covalently bonded to the CMN surface via EDC/NHS-mediated activation of its carboxyl group, enabling amide bond formation with primary amine groups of CMN.^32^ Briefly, Ga (20 mg/ml) was prepared in 100% ethanol, and a final Ga concentration of 140 μg/ml (298 μM) was activated with EDC (26.1 mM) and NHS (43.4 mM) in PBS at room temperature for 30 minutes. The activated Ga solution was then added to 1 ml of ghost RBCs containing 5.6 × 10^7^ ± 0.6 × 10^7^ cells/ml in PBS, and the mixture was gently stirred for 3 hours at room temperature to facilitate amide bond formation. The functionalized RBCs were pelleted by centrifugation (14,000 × g, 10 minutes), washed twice with PBS to remove unbound reagents, and subsequently used to prepare the Ga-functionalized CMN.

#### La-modified RBCs

Lactobionic acid (La) was covalently bonded to the CMN surface using EDC/NHS coupling, where the carboxyl groups of La were activated and subsequently coupled to the amine groups on the CMN surface to form stable amide bonds.^29^ As explained above for Ga-functionalized RBCs, the carboxyl group of La was first activated with EDC and NHS, then reacted with ghost RBCs to prepare the functionalized CMN. Briefly, La (5.0 mg, 100 mM) was dissolved in 140 µL of PBS, followed by the addition of EDC (13.4 mg, 700 µL of 100 mM EDC in PBS) and NHS (8.1 mg, 700 µL of 100 mM NHS in PBS). The reaction mixture was stirred for 30 minutes at room temperature to generate the La-NHS ester intermediate. The activated La solution was added dropwise to 1 ml of RBC suspension under gentle stirring, and the mixture was incubated for 3 hours at room temperature to allow covalent amide bond formation between La and the free amine groups on the RBC membrane surface. Lastly, the unreacted La was removed by centrifugation (14,000 × g, 10 min), and washed twice with PBS, and used to prepare the La-functionalized CMN.

#### Lf-modified RBCs

Lactoferrin (Lf) was covalently bonded onto the RBC membrane through EDC/NHS chemistry, where the carboxyl groups of Lf reacted with primary amine residues of RBC membrane proteins to form stable amide bonds.^34,35^ Briefly, Lf (2 mg/ml) was mixed with 26.1 mM EDC and 43.4 mM NHS dissolved in 1 ml of PBS, and the reaction mixture was stirred for 30 minutes at room temperature to activate the carboxyl groups of Lf. The activated Lf solution was then added dropwise to the RBC suspension, and the mixture was stirred gently for 3 hours at room temperature to allow covalent coupling between the amino groups of membrane proteins and the activated Lf. Finally, the modified RBCs were washed with PBS to remove unbound Lf and residual reagents.

Finally, Ga-, La-, and Lf-modified RBCs were passed through an extruder system equipped with 3, 0.8, and, lastly, 0.2 μm filters to obtain uniform, nanosized particles with a size range below 200 nm.

### 2.3. Physicochemical characterization of hepatocyte-targeted CMN

Nanoparticle tracking analysis (NTA) and transmission electron microscopy (TEM) were employed to determine the shape and size of the CMN and hepatocyte-targeting CMN, including CMN-Ga, CMN-La, and CMN-Lf. For TEM imaging, a 30 μl drop of freshly prepared CMN was placed on a 200-mesh carbon-coated copper grid and allowed to settle for 5 minutes. The grid was then washed twice with ultrapure DI water by gentle dipping, and excess liquid was blotted with filter paper. Subsequently, the grids were negatively stained with 0.5% uranyl acetate, air-dried, and imaged using a JEOL JEM-1230 transmission electron microscope. Particle size was further confirmed using NTA on a ZetaView Analyzer (Particle Metrix Inc.), and zeta potential values were also measured. After confirming nanoparticle size, the functionalized CMN samples (CMN-Ga, CMN-La, and CMN-Lf) were characterized by Fourier transform infrared spectroscopy (FTIR) using a Nicolet Summit LITE FTIR Spectrometer (Fisher Scientific). Protein concentrations of RBCs and CMN-based samples were determined using the Pierce BCA Protein Assay Kit, following the manufacturer’s protocol.

### 2.4. Cytocompatibility of hepatocyte-targeted CMN

The cytocompatibility of CMN and hepatocyte-targeted CMN functionalized with Ga, La, and Lf were assessed using the LIVE/DEAD^™^ Cell Imaging Kit (Thermo Fisher, R37601) and MTS assay (Promega, G3580). HepG2 cells were seeded at 50,000 cells per well in 48-well plates and incubated for 48 hours at 37°C in a 5% CO_2_ incubator. After replacing the medium with fresh culture media, cells were treated with CMN, CMN-Ga, CMN-La, or CMN-Lf (∼4.5 × 10^9^ particles/well) and incubated for 24 hours.

Live/Dead staining was then performed according to the manufacturer’s protocol, and fluorescence images were acquired using a Nikon Eclipse Ti2-E microscope (Nikon Instruments Inc., Canada). For the MTS assay (to assess mitochondrial activity), the treatment medium was discarded, and 225 μL of fresh media with 25 μL of MTS reagent was added to each well. Following a 3-hour incubation, absorbance was measured at 490 nm using a SPARK multimode microplate reader (Tecan, USA).

We further confirmed whether CMN induces cell inflammation by seeding THP-1 cells at 1 × 10^6^ cells per well in a 24-well plate. THP-1 cells were first differentiated into macrophages by stimulation with 50 ng/mL phorbol 12-myristate 13-acetate (PMA; Sigma, P8139) for 48 hours. An additional 24-hour incubation in fresh medium was then performed to remove residual PMA and non-adherent cells. The macrophages were treated with CMN (∼4.5 × 10^9^ particles per well), lipopolysaccharide (LPS; Sigma, L6929) as a positive control, or left untreated for 24 hours. TNF-α and IL-6 secretion by macrophages was quantified using a Quantikine^™^ ELISA kit (R&D Systems) according to the manufacturer’s protocol.

### 2.5. Hepatocyte targeting ability of functionalized CMN

To identify the optimal hepatocyte-targeting CMN, we conducted cellular uptake assays in three different cell types, including HUVECs (Human Umbilical Vein Endothelial Cells, passage 7), hSCs (Human Stellate Cells, passage 6), and HepG2 (hepatocytes, passage 29) cells.^32,34^ Cells were seeded at a density of 50,000 cells/well in a 24-well plate and incubated for 48 hours. CMN and the functionalized CMN (CMN-Ga, CMN-La, and CMN-Lf) were tagged with the fluorescent lipophilic membrane dye, DiI (Millipore Sigma, 42364).^21,36^ A stock solution of DiI at a concentration of 1 mg/ml was diluted at a ratio of 1:100 and mixed with an equal volume of CMN or functionalized CMN. After incubation at 4°C for 1 hour, samples were loaded into an Amicon^®^ Ultra Centrifugal Filter (10 kDa MWCO, 0.5 ml volume) and centrifuged at 14,000g for 8 min to remove free dye molecules. The centrifugation step was repeated three times, after which labelled CMN were resuspended in DPBS, quantified using NTA, and later prepared for the cell internalisation experiment.

An average of 2.68 × 10^9^ particles/ml of CMN, CMN-Ga, CMN-La, and CMN-Lf were added to the cells, followed by incubation for 2 hours. Cells were then thoroughly washed with PBS, trypsinized, and resuspended in 200 µl PBS. Finally, samples were measured using a CytoFlex flow cytometer (Beckman Coulter, Canada).

The distribution of CMN and CMN-Lf (identified as the optimal preparation for liver targeting) in hepatocytes was further qualitatively analysed by fluorescent microscopy. After treatment with the nanoparticles, cells were washed and then fixed with 4% paraformaldehyde for 10 minutes, followed by DPBS washes. Next, cells were stained with phalloidin-iFluor 488 (ab176753, Abcam) for actin staining and counterstained with DAPI (Fisher Scientific, 5087410001) according to the manufacturer’s protocol. After staining, mounting medium was added to the wells, and coverslips were placed on the wells before taking the fluorescent images using a Nikon Eclipse Ti2-E microscope (Nikon Instruments Inc., Canada). Images were analysed using an image processing software (Imaris, Oxford Instruments, USA).

### 2.6. Endocytic pathway analysis of CMN and CMN-Lf

After identifying the most effective hepatocyte-targeting CMN, the endocytic pathways involved in the cellular uptake of CMN and CMN-Lf were investigated.^37^ HepG2 cells were seeded at the same density as described above in 24-well plates. Before nanoparticle treatment, cells were pre-treated with specific pharmacological inhibitors to block endocytosis pathways: chlorpromazine (20 μg/mL; Millipore Sigma, C-904), nystatin (100 U/mL; Millipore Sigma, N9150), and amiloride (133 μg/mL; Millipore Sigma, 129876). Cells were incubated with these inhibitors for 1 hour at 37°C.

Following pre-treatment, DiI-labelled CMN and CMN-Lf at a concentration of 2.65 × 10^9^ particles/ml were added to the cells and incubated for 2 hours. After incubation, media were removed, cells were washed three times with DPBS, trypsinized, resuspended in PBS, and immediately analyzed by flow cytometry. Cellular uptake of nanoparticles was calculated using the following equation. Finally, cellular uptake of nanoparticles, with and without inhibitors, was compared to identify the specific endocytic pathways responsible for CMN and CMN-Lf internalization.

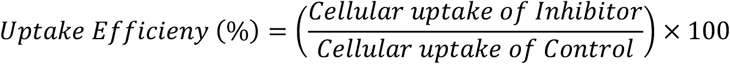

### 2.7. Cell apoptosis analysis

We further confirmed that CMN and CMN-Lf were non-cytotoxic by assessing apoptosis, using the Annexin V and PI-based Cell Apoptosis Kit from Biotium (30061). In brief, HepG2 cells were seeded at 2 × 10^5^ cells/ml in 12-well plates and incubated for 24 hours. Cells were then treated with CMN or CMN-Lf (1.9 × 10^10^ particles/ml) and incubated for an additional 24 hours. A positive control was prepared by incubating the cells at 55°C for 20 min (heat shock), followed by an additional 24 hours before the cell apoptosis assay.^21^ After treatment, media were removed, and cells were prepared for cell apoptosis analysis according to the manufacturer’s protocol. All samples were analysed using a flow cytometer, and the resulting data were assessed using FlowJo analysis software.

### 2.8. *In Vitro* oleic and palmitic acid-induced hepatocyte steatosis model and treatment

To confirm the bioactivity of CMN-Lf, we first loaded the model drug Resmetirom (Res), which has recently been approved for the treatment of MASLD. The drug loading process was based on our previously established protocol.^36^ Briefly, after passing the Lf-conjugated RBC fragments through a 0.8 μm filter, we mixed the drug (Res) into 1 ml of the RBC suspension, achieving a 1 mM Res concentration. Then, we extruded the mixture through a 0.2 μm filter twice, removed the unloaded drug using an Amicon^®^ Ultra Centrifugal Filter (10 kDa MWCO, 0.5 ml volume), and centrifuged at 14,000g for 8 minutes. Next, we collected the filtered solution and measured the absorbance at 298 nm using a TECAN Spark^®^ multimode microplate reader. Finally, we calculated the drug loading efficiency of Res-loaded CMN-Lf (CMN^Lf^-Res) using the following equations:

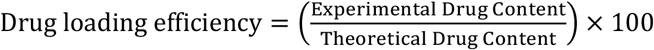

Separately, we prepared an *in vitro* hepatocyte steatosis model to assess the efficacy of CMN^Lf^-Res. To do so, we exposed HepG2 cells to a combination of oleic acid and palmitic acid.^21^ Growth medium (EMEM containing 10% FBS, 1% Pen-Strep, and 1% L-glutamine) was supplemented with palmitic acid and oleic acid (2:3 palmitic acid to oleic acid ratio; 2:1 fatty acid to BSA ratio), achieving a total fatty-acid concentration of 1 mM. Cells were seeded at a density of 2.5 × 10^5^ into each well of a 12-well plate and incubated for 24 hours, followed by overnight incubation with fatty acid-containing medium. The next day, the medium was replaced with basal growth medium, and the cells were treated with Res (100 μM), CMN-Lf, or CMN^Lf^-Res (100 μM) for 24 hours.^38^

After treatment, Oil Red O staining, triglyceride assays, and ALT and AST activity measurements were performed. Neutral lipids were determined using the lipophilic stain, Oil Red O (ORO). Fresh 0.5% (w/v) ORO solution was prepared following our published protocol.^21^ Cells were washed with DPBS and then fixed with 4% paraformaldehyde for 10 minutes. ORO solution was added to the cells, followed by incubation at room temperature for 15 minutes. Finally, the cells were thoroughly washed with DPBS before brightfield images were captured using a Nikon Eclipse Ti2-E microscope. Triglycerides, ALT, and AST activities were measured according to the manufacturer’s protocols.

### 2.9. Statistical analysis

Experiments were performed in triplicate (three biological replicates), and data are presented as mean ± standard deviation. Statistical differences between groups were evaluated using one-way or two-way ANOVA followed by Tukey’s post hoc test, performed in GraphPad Prism. A p-value < 0.05 was considered statistically significant (*p < 0.05, **p < 0.01, ***p < 0.001, ****p < 0.0001).

## 3. Results and Discussions

### 3.1. Physicochemical confirmation of CMN-Ga, CMN-La, and CMN-Lf functionalization

The morphology of CMN prepared from mouse RBCs was examined by TEM, which confirmed the intact, spherical structure of the nanoparticles (**Figure 2a**). Particle size and zeta potential of CMN, CMN-Ga, CMN-La, and CMN-Lf were assessed using NTA (**Figures 2b and 2c**). CMN-Ga exhibited a noticeable increase in size compared to unmodified CMN. The average sizes of CMN, CMN-Ga, CMN-La, and CMN-Lf were 154.6 ± 6.7 nm, 207.7 ± 13.6 nm, 159.5 ± 7.2 nm, and 165.7 ± 6.8 nm, respectively. Significant changes in zeta potential were observed for CMN-Ga and CMN-Lf. Because lactoferrin is a positively charged glycoprotein, its conjugation decreased the zeta potential of CMN-Lf, confirming functionalization. Moreover, we observed a significant increase in CMN-Lf protein concentration compared to CMN, confirming the presence of Lf in CMN (**Figure S1**). FTIR analysis further validated functionalization. FTIR spectra (**Figure 2d**) showed the characteristic amide band at 1640 cm^-1^ in CMN-Ga and CMN-La, along with the disappearance of the carboxyl-group bands of Ga (1705 cm^-1^) and La (1740 cm^-1^). CMN-Lf also displayed a prominent amide band at 1640 cm^-1^, confirming the functionalization of Lf into CMN. Taken together, these physicochemical characterization data confirm the successful functionalization of CMN with Ga, La, and Lf using EDC/NHS chemistry.

**Figure 2.**
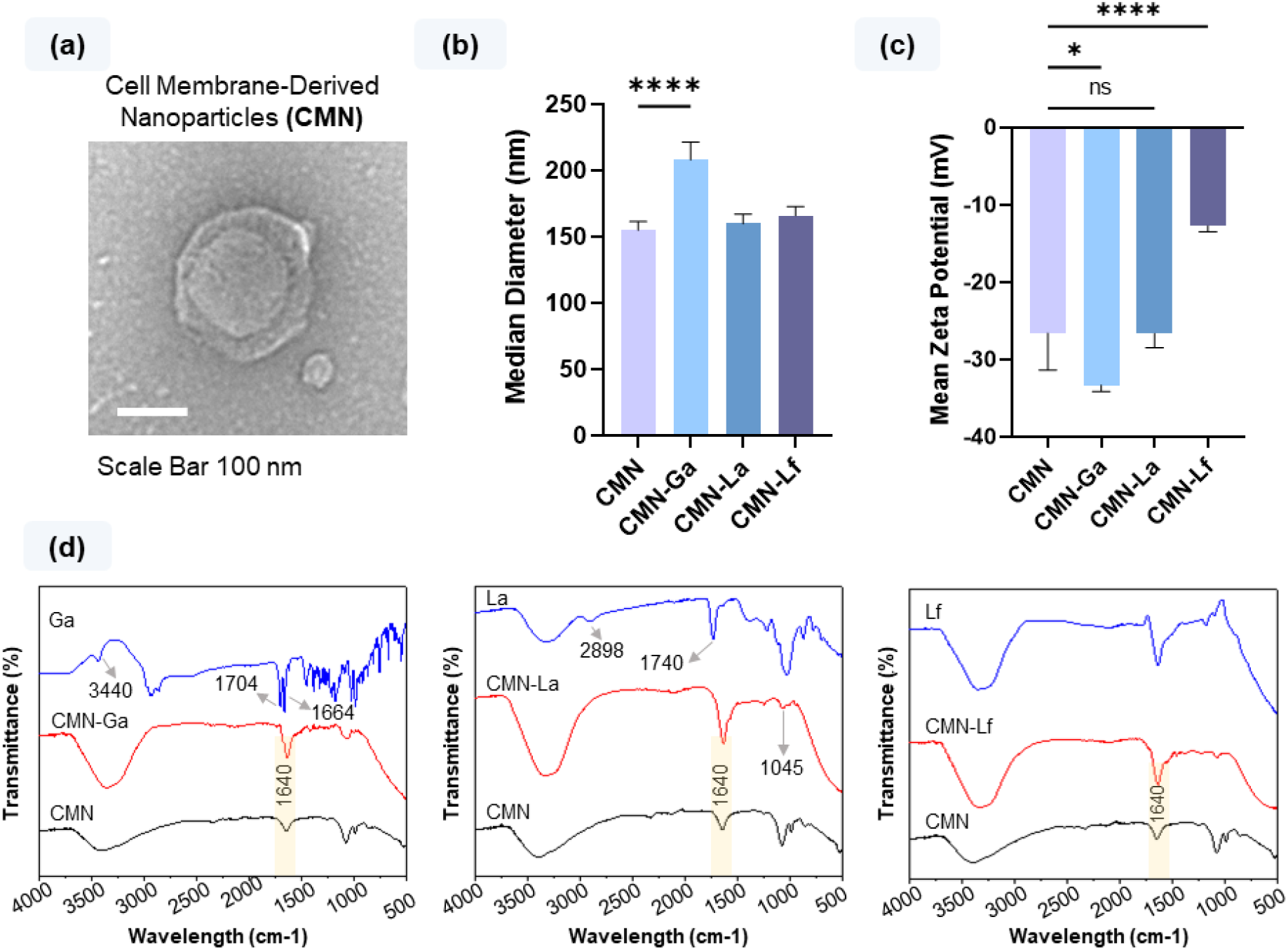
Characterization of hepatocyte-targeting CMN. a) TEM image displays the shape and morphology of CMN prepared using the cell extrusion method. b) Median diameters of CMN and surface-modified CMNs, including CMN-Ga, CMN-La, and CMN-Lf, were determined using NTA, confirming the nano-size of the particles. c) Zeta potential values of CMN-Ga and CMN-Lf, confirming the successful conjugation of Ga and Lf. d) FTIR analysis further corroborates the functionalization of Ga, La, and Lf to CMN. Results are shown as means ± SD. ****p < 0.0001, *p < 0.05, ns = non-significant. Scale bar = 100 nm.

### 3.2. *In Vitro* Cytocompatibility of CMN-Ga, CMN-La, and CMN-Lf in hepatocytes

Next, we evaluated the cytocompatibility of functionalized CMN *in vitro* using live/dead staining and an MTS assay. **Figure 3a** presents phase contrast and fluorescence images of HepG2 cells treated with CMN, CMN-Ga, CMN-La, and CMN-Lf, showing predominantly live cells (Calcein-AM, green) with minimal dead-cell staining (BOBO-3, red), indicating negligible cytotoxicity for all formulations. Consistent with these qualitative results, the MTS assay (**Figure 3b**) showed no significant differences in mitochondrial activity or overall viability among CMN (control), CMN-Ga, CMN-La, and CMN-Lf. Furthermore, the inflammatory activity of CMN was investigated using THP-1–differentiated macrophages. After 24 hours of incubation, CMN-treated macrophages exhibited no significant increase in inflammatory cytokine secretion, including TNF-α and IL-6, as shown in **Figure S2**. These findings confirm that functionalized RBC-derived CMN are cytocompatible and suitable for subsequent hepatocyte-targeting studies.

**Figure 3.**
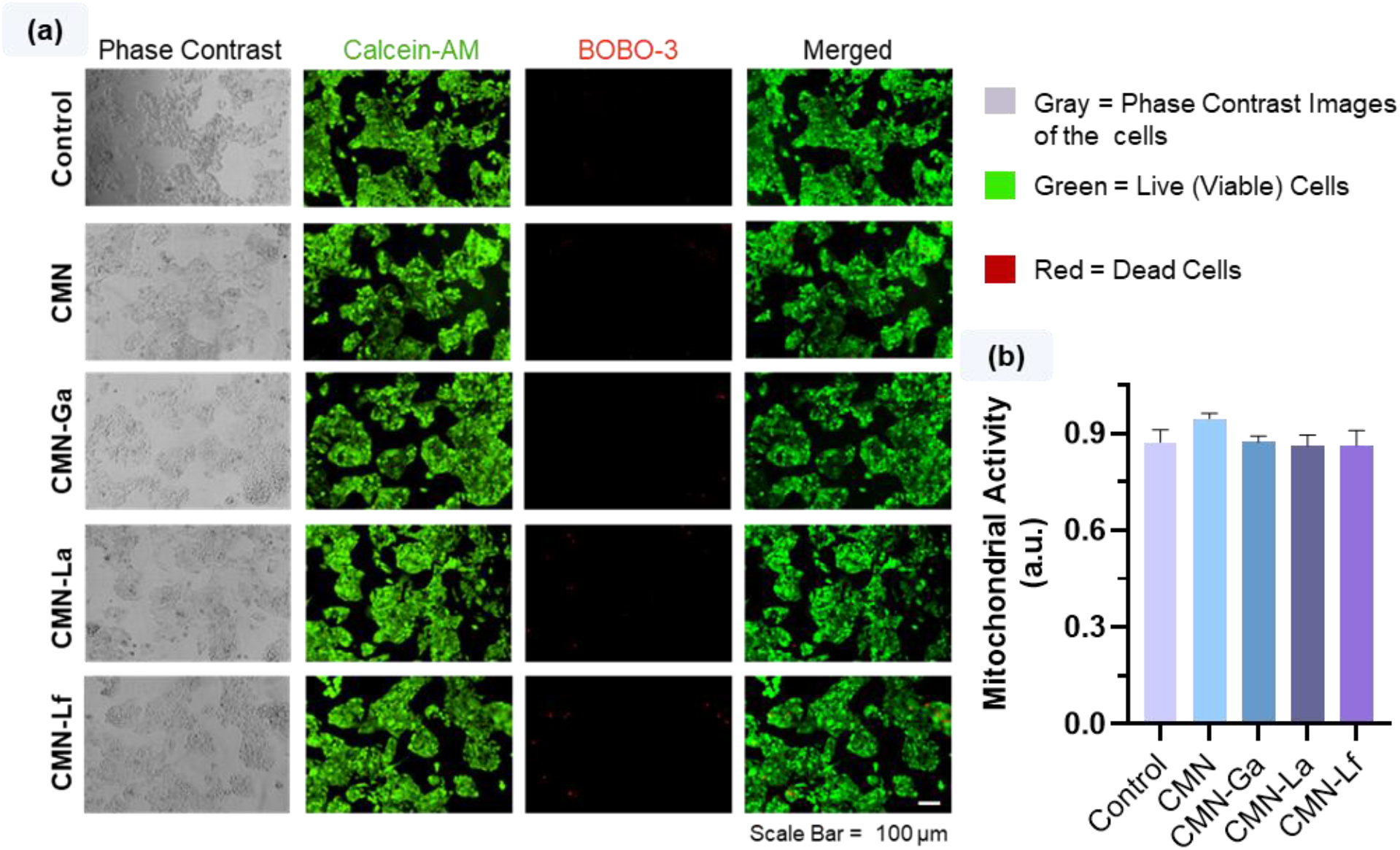
Cytocompatibility of hepatocyte-targeting CMN in HepG2 cells.. a) Qualitative live (green, Calcein-AM) and dead (red, BOBO-3) images of hepatocytes treated with CMN, CMN-Ga, CMN-La, and CMN-Lf, showing no observable cytotoxicity between the groups. b) MTS assay results showed no changes in mitochondrial activity, further confirming the cytocompatibility of all CMN formulations. Results are shown as means ± SD. Scale bar = 100 µm.

### 3.3. Cellular uptake analysis of CMN-Ga, CMN-La, and CMN-Lf for hepatocyte-specific targeting efficiency

Targeted delivery to hepatocytes relies on receptor-mediated interactions, and ligands such as Ga, La, and Lf have been widely explored to enhance nanoparticle recognition by liver cells.^32,39,40^ To evaluate the effect of these ligands on cell-specific uptake, we quantified the internalization of CMN, CMN-Ga, CMN-La, and CMN-Lf in endothelial cells, stellate cells, and hepatocytes using flow cytometry. DiI-labeled functionalized nanoparticles were prepared following the established protocol, and the results are shown in **Figure 4**. In endothelial cells, CMN, CMN-Ga, and CMN-La exhibited comparable internalization, while CMN-Lf showed a slight decrease. Since endothelial cells do not express hepatocyte-associated receptors, such as ASGPR, this similarity in nanoparticle uptake was expected and suggests that none of the ligands promote unintended endothelial targeting.^41^ The slightly reduced uptake of CMN-Lf may reflect lower nonspecific association due to the surface characteristics introduced by lactoferrin, as previously reported for Lf-functionalized nanoparticles.^34^ In stellate cells, CMN, CMN-Ga, and CMN-La showed low, similar uptake, whereas CMN-Lf showed an apparent increase in internalization. However, overall uptake was lower than in the hepatocytes (HepG2 cells), suggesting a lack of unintended binding to non-parenchymal liver cells. Finally, in hepatocytes, CMN-Lf showed a significant 3.4-fold increase in uptake relative to CMN. This observation is consistent with previous reports indicating that hepatocytes efficiently recognize lactoferrin-modified nanocarriers, due to high expression of ASGPR on the hepatocyte surface.^34,39^ Interestingly, a single HepG2 cell expresses around 76,000 ASGPR receptors, which likely contributes to the enhanced interaction of CMN-Lf with hepatocytes compared with non-target cells.^30^ Conversely, uptake of CMN-Ga and CMN-La was not significantly increased in HepG2 cells compared to CMN alone. Previous studies have shown that both Ga-mediated targeting and galactose-mediated ASGPR uptake heavily depend on ligand density on the nanoparticle surface^30,42^, and CMN surface structure may partially shield Ga or La molecules, limiting ligand-receptor binding to hepatocytes. Based on these concepts, our data suggest that Ga and La may require higher density or improved display orientation to achieve effective hepatocyte targeting when conjugated to CMN. Overall, our findings indicate that CMN-Lf provides the most effective and most selective hepatocyte targeting.

**Figure 4.**
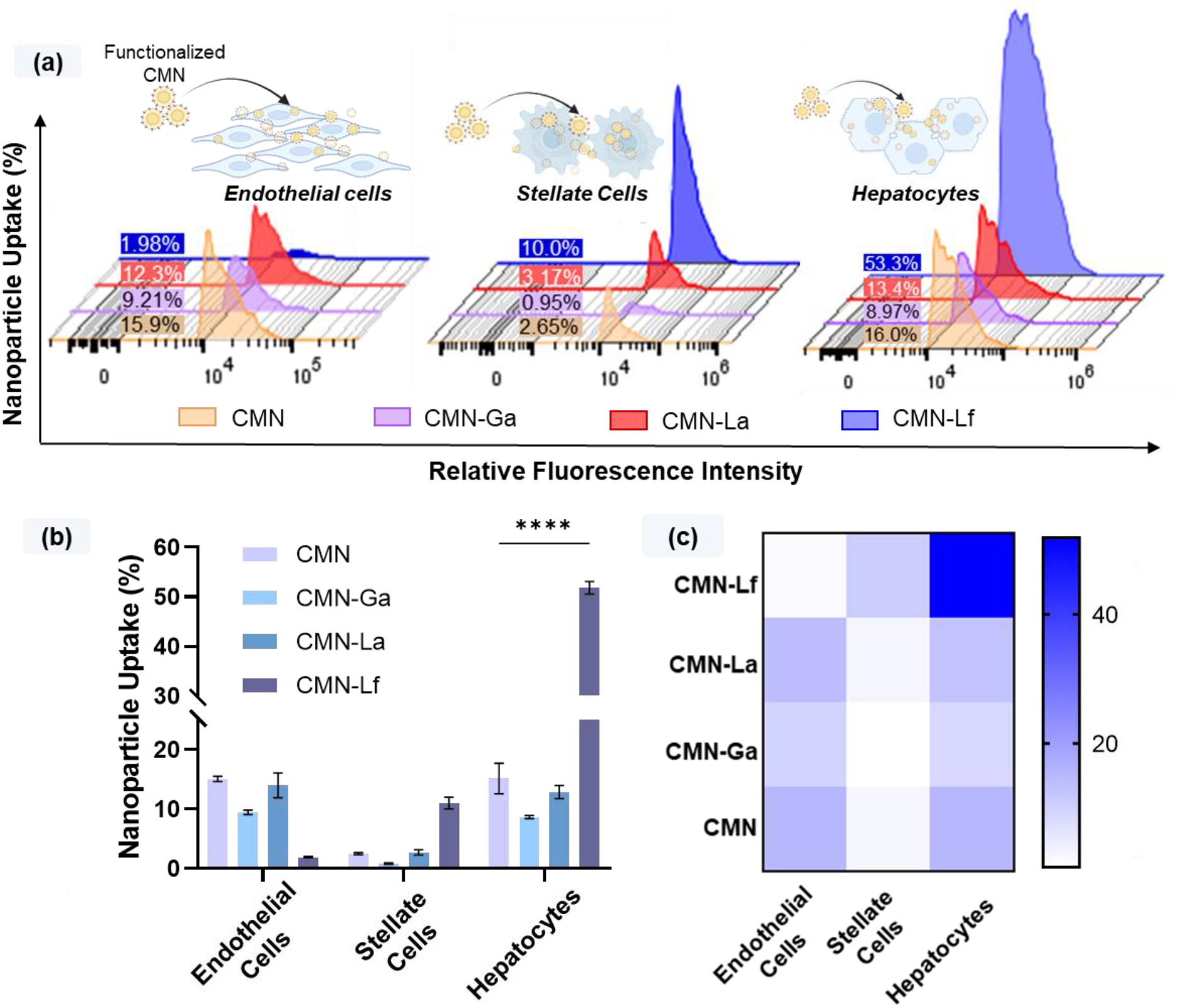
Screening the cellular uptake efficiency of CMN-Ga, CMN-La, and CMN-Lf in endothelial cells, stellate cells, and hepatocytes. a) Flow cytometry staggered histograms showing the representative cellular uptake of CMN, CMN-Ga, CMN-La, and CMN-Lf. b-c) Quantitative analysis presented as a bar graph and heat map, confirming that CMN-Lf exhibits the highest hepatocyte-specific targeting efficiency. In contrast, CMN-Ga and CMN-La did not show a significant increase in hepatocyte uptake. Results are expressed as mean ± SD. ****p < 0.0001.

### 3.4. Mechanism of cellular uptake of CMN-Lf by hepatocytes

Prior to investigating uptake mechanisms, we visually confirmed cellular uptake of CMN and CMN-Lf by fluorescence cytochemistry. Acquired images (**Figure S3)** were deconvolved to improve clarity, as shown in **Figure 5a**. Hepatocytes treated with CMN exhibited a limited number of internalized nanoparticles, visible as red dots. In contrast, CMN-Lf showed markedly greater cytosolic nanoparticle accumulation, as indicated by arrows. We next aimed to identify which endocytic pathway (clathrin-mediated, caveolae-mediated, or micropinocytosis) mediated CMN and CMN-Lf internalization. For this purpose, we used pharmacological inhibitors, including chlorpromazine to inhibit clathrin-mediated uptake, nystatin to inhibit caveolae-mediated uptake, and amiloride to inhibit micropinocytosis. As shown in **Figure 5b**, all three inhibitors significantly reduced uptake of both CMN and CMN-Lf. For CMN, the caveolae-mediated pathway contributed most to internalization, consistent with previous findings that EV- and liposome-based nanocarriers primarily enter hepatocytes via caveolae-mediated endocytosis.^43,44^ One study further showed that pre-treating hepatocytes with methyl-β-cyclodextrin, an inhibitor of caveolae-mediated uptake, reduced lipid-based nanoparticle internalization by 60%, suggesting caveolae-mediated internalization.^45^ For CMN-Lf, we observed the most significant reduction in uptake when clathrin-mediated endocytosis was inhibited, with smaller decreases observed upon inhibition of caveolae and micropinocytosis pathways. These data are consistent with the known binding of Lf to hepatic lectin-1 (HL-1), which is a major subunit of ASGPR.^46^ Upon ligand binding, ASGPR clusters into clathrin-coated pits, where clathrin and adaptor proteins drive vesicle formation and internalization. After entry, the particle sheds its coat and matures through endosomal compartments, where acidification releases the ligand. ASGPR is then recycled back to the hepatocyte surface, allowing repeated rounds of efficient uptake.^30,37,47^ Thus, our results indicate that CMN-Lf is largely internalized via the clathrin-mediated pathway, consistent with clathrin-dependent endocytosis of ASGPR-targeting ligands.

**Figure 5.**
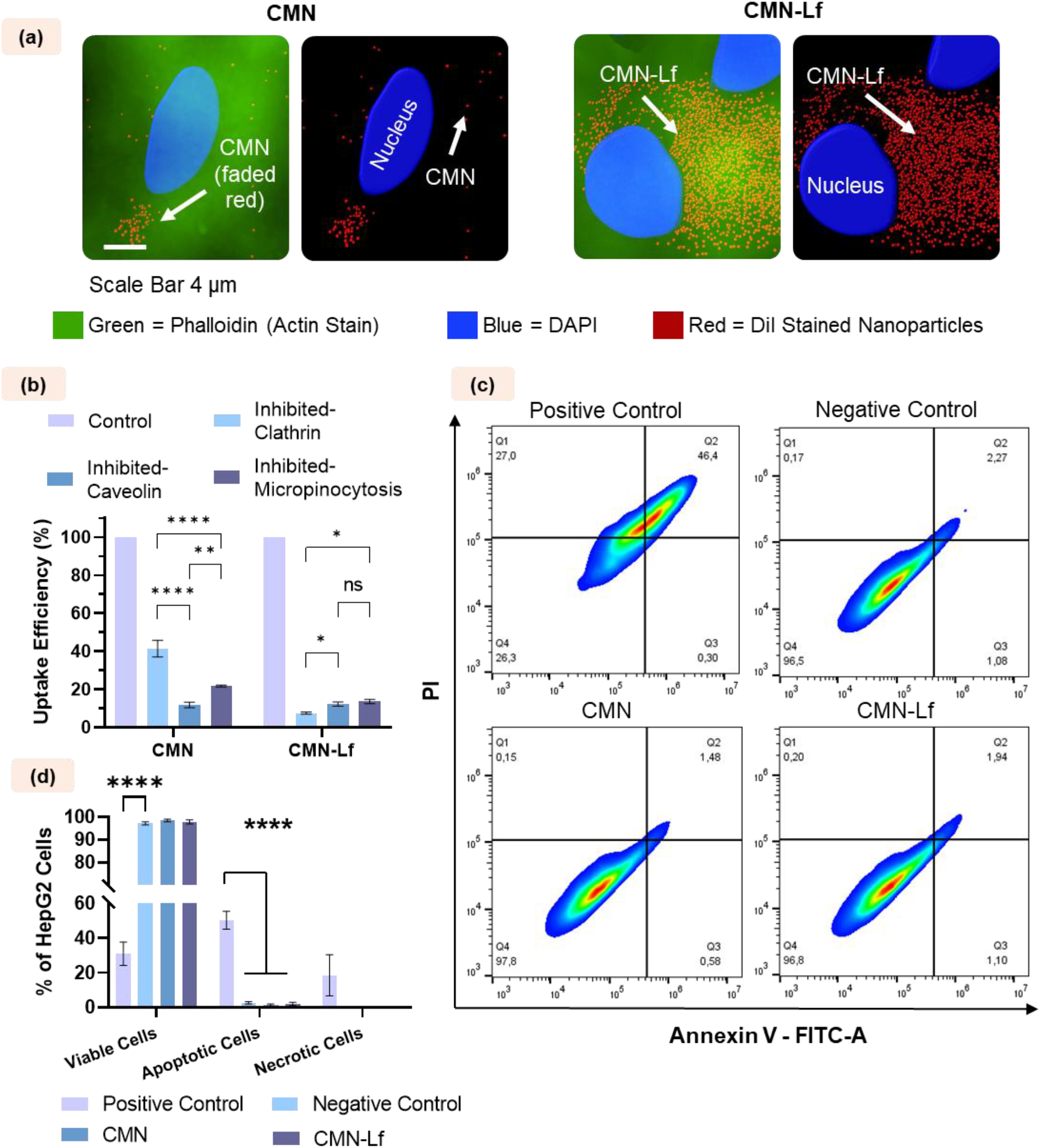
Visualization of CMN-Lf uptake, identification of internalization pathways, and assessment of apoptosis in hepatocytes. a) Processed fluorescence images show that CMN-Lf (red dots, indicated by white arrows) accumulates more prominently around the nuclei, confirming higher cellular uptake compared with CMN. b) Receptor-mediated endocytosis pathway studies indicate that CMN is primarily internalized through the caveolin-mediated pathway, whereas CMN-Lf shows significant dependence on clathrin-mediated uptake. This observation aligns with reports demonstrating that ASGPR-targeting ligands are predominantly internalized via clathrin-mediated endocytosis. c-d) Apoptosis analysis confirms that CMN-Lf does not affect cell viability and shows no significant increase in apoptosis compared with the negative control and CMN groups. Scale bar 4 μm. Results are expressed as mean ± SD. ****p < 0.0001, **p < 0.01, *p < 0.05, ns = non-significant.

Subsequently, we further evaluated the cytocompatibility of CMN-Lf by assessing apoptosis. **Figure 5c** shows representative flow cytometry quadrant plots for control, CMN, and CMN-Lf treated HepG2 cells, with the quantitative analysis summarized in **Figure 5d**. There were no significant differences in viable cells among the negative control, CMN, and CMN-Lf-treated cells. Similarly, CMN-Lf did not induce apoptosis compared with the control. These findings further confirm that CMN-Lf can be a safe nanotherapeutic platform for drug delivery to hepatocytes.

### 3.5. Bioactivity of Res-loaded CMN-Lf in an *in vitro* steatotic hepatocyte model

To determine whether our CMN-Lf nanodelivery platform could improve the efficacy of an established MASLD drug that is known to decrease hepatocyte steatosis, we exposed lipid-loaded HepG2 cells (a model of hepatocyte steatosis) to CMN-Lf containing the thyroid hormone receptor β (THR-β) agonist, resmetirom (Res). Once inside the cell, Res binds to THR-β and mimics the transcriptional activity of free triiodothyronine (FT3), as illustrated in **Figure 6a**. This interaction facilitates heterodimerization of THR-β with the retinoid X receptor (RXR), enabling the complex to bind thyroid hormone response elements (TREs) on specific target genes. ^12^ The activation of this signaling pathway induces key metabolic pathways involved in hepatic lipid homeostasis, including enhanced cholesterol catabolism, lipophagy stimulation, and increased fatty acid β-oxidation.^48^

**Figure 6.**
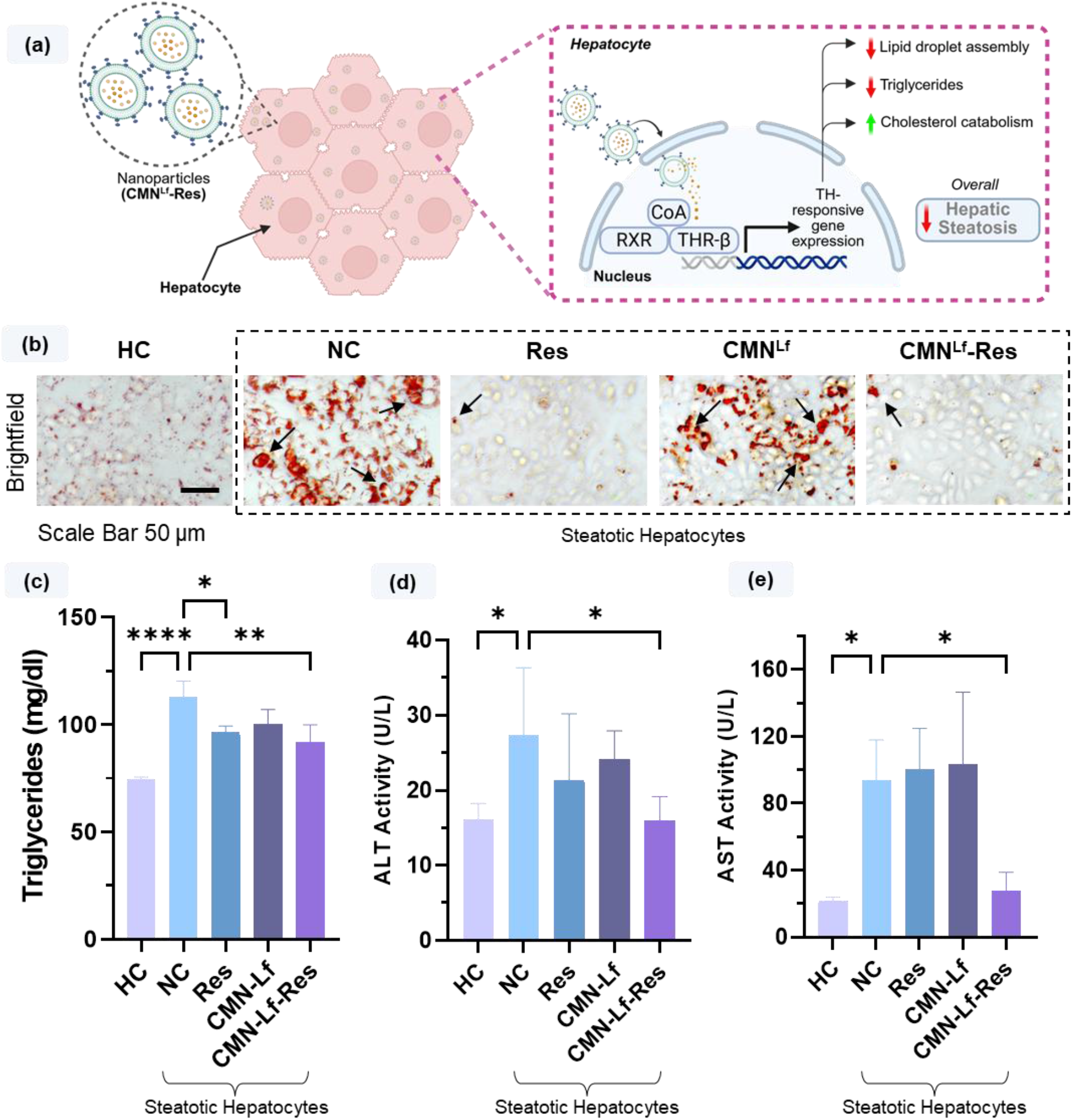
Evaluating the efficacy of resmetirom (Res)-loaded CMN-Lf in a hepatocyte steatosis model. a) The schematic illustrates the mechanism of Res upon release from CMN-Lf in the cytosol, leading to reduced triglycerides and hepatic lipids. b) Oil Red O staining suggests a marked reduction in neutral lipid accumulation in the CMN^Lf^-Res treated group compared with the negative control (NC). c-e) Quantitative analysis shows significant decreases in intracellular triglycerides, ALT levels, and AST activity in CMN-Lf-Res treated cells relative to NC. Scale bar = 50 μm. Results are expressed as mean ± SD. **p < 0.01, *p < 0.05.

Following our established protocol for drug loading^21,36^ we observed that 59.33 ± 3.5% Res was successfully loaded into CMN-Lf. Further, the drug-loaded CMN-Lf can be visualized by the TEM images provided in **Figure S4**. We treated steatotic HepG2 cells with Res-loaded CMN-Lf and assessed neutral lipid accumulation by ORO staining (**Figure 6b**). These data suggest that the Res-loaded CMN-Lf markedly decreased neutral cytosolic lipid droplets compared to the NC (Negative Control, hepatocyte steatosis) and CMN-Lf. We next quantified triglycerides, ALT, and AST biochemically, using commercial kits. **Figure 6c** shows that Res-loaded CMN-Lf significantly reduced triglycerides compared to NC, but no significant difference in the triglyceride level between Res and CMN^Lf^-Res treated cells. However, we observed significant reductions in ALT and AST levels in CMN^Lf^-Res-treated cells, but not in Res-treated cells, compared with NC. These data suggest enhanced efficacy of CMN^Lf^-Res in decreasing lipid-induced cell damage, as indicated by diminished release of these hepatocellular enzymes. By combining hepatocyte-targeted uptake with a clinically relevant MASLD drug, CMN-Lf may have the potential to enhance therapeutic response and overall disease improvement. This nanodelivery strategy may be particularly useful for the large number of patients with MASH who do not respond to Res treatment (approximately 3 in 4 individuals).^49^ Targeted nanocarriers, such as CMN-Lf, may help address this gap by enhancing hepatocyte-specific drug accumulation and potentially increasing the efficacy of such therapeutic agents.

## 4. Conclusion

Herein, we designed a hepatocyte-targeting Res-loaded CMN to enhance therapeutic efficacy and help mitigate hepatocyte steatosis. We functionalized CMN with three different targeting ligands, including Ga, La, and Lf, and confirmed their functionalization by size, zeta potential, and FTIR analyses. Subsequently, we performed a thorough *in vitro* evaluation of their targeting potential in endothelial, stellate cells, and hepatocytes. Our results showed that CMN-Lf outperformed CMN-Ga and CMN-La in terms of hepatocyte-specific internalization. We identified the cellular internalization mechanisms of selected hepatocyte-targeting CMN-Lf, showing that for CMN and CMN-Lf, caveolae and clathrin, respectively, are responsible for cellular uptake. This aligns with previous reports that ASGPR-targeting ligands are internalized by the clathrin-mediated endocytosis pathway. In contrast, lipid-based nanoparticles, such as CMN, are internalized via the caveolin-mediated endocytosis pathway. Finally, and most importantly, we show that the bioactivity of highly specific hepatocyte-targeting CMN-Lf loaded with Res has the potential to improve efficacy in an *in vitro* hepatocyte steatosis model. Our results confirm that CMN^Lf^-Res significantly reduces intracellular triglycerides, although not to a greater extent than Res alone. But, remarkably, CMN^Lf^-Res attenuates liver enzyme (ALT and AST) release more effectively than Res alone. Overall, our study demonstrates that the CMN-Lf may have the potential to enhance the therapeutic efficacy of Res in alleviating liver steatosis. In future studies, we plan to investigate its hepatocyte targeting ability and enhanced therapeutic potential in an *in vivo* MASLD model.

## Acknowledgments

Arghya Paul is thankful to the following funding agencies for providing support: New Frontiers in Research Fund (NFRF)-Exploration Stream (NFRFE-2024-01126), Early Research Award (ERA) from the Province of Ontario, Canada Research Chairs Program of the Natural Sciences and Engineering Research Council (NSERC) of Canada (CRC-2024-00196), Wolfe-Western Fellowship At-Large for Outstanding Newly Recruited Research Scholar, and Canadian Institutes of Health Research Operating Grant (CIHR-IMHA, Grant no: 185629). Alap Ali Zahid would like to acknowledge the support from The Ivan Malek Scholarship and The E.G.D. Murray Scholarship in Biochemical Engineering, The University of Western Ontario, Canada. The authors would also like to thank BioRender, as some images and illustrations were created with BioRender.com.

