## Supplementary File for "Surface Functionalized RBC Membrane-Derived Nanoparticles for Targeted Drug Delivery to Attenuate Fatty Liver Disease"

**Keywords**

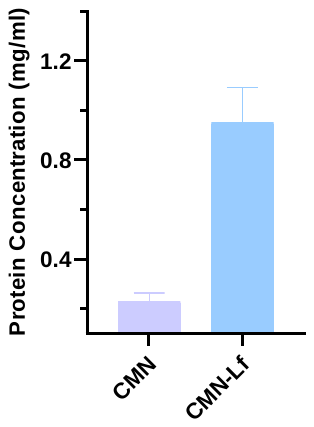

**Figure S1**. Protein concentration of CMN and CMN-Lf. CMN-Lf showed a significant increase in protein concentration compared to its CMN alone, confirming the presence and functionalization of Lf on CMN (n=3).

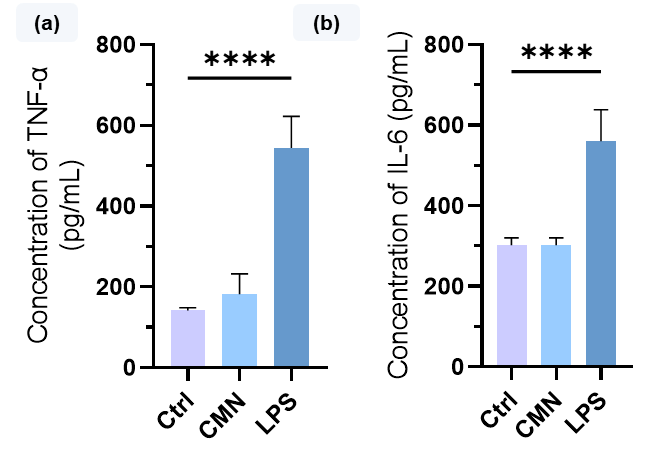

**Figure S2.** The graph shows ELISA analyses of the inflammatory cytokines TNF-α and IL-6 secreted by THP-1 differentiated macrophages after 24 h stimulation with CMN and LPS (positive control) or left untreated. No significant inflammatory response was observed for CMN (*n* = 3).

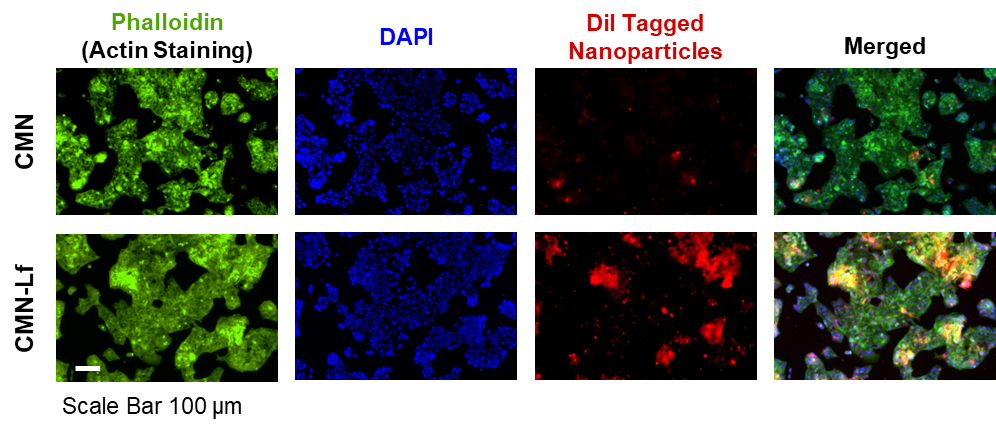

**Figure S3.** Fluorescent images before deconvolution for studying the cellular uptake analysis of CMN and CMN-Lf by hepatocytes. CMN and CMN-Lf were tagged with a fluorescent label dye DiI. The CMN-Lf group demonstrates a significant increase in red fluorescence, confirming higher nanoparticle uptake by hepatocytes.

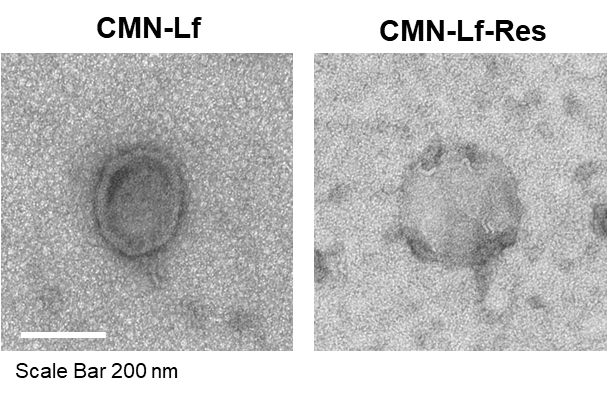

**Figure S4**. TEM images of CMN-Lf and CMN-Lf-Res.
